# Mechanism underlying PilC/PilY1 role in type 4a pilus biogenesis at a late stage occurring after filament assembly

**DOI:** 10.64898/2026.09.18.752573

**Authors:** Milica Denic, Morgane Wuckelt, Véronique Roig-Zamboni, Odile Valette, Nicolas Biais, Latifa Elantak, Gerlind Sulzenbacher, Mathieu Coureuil, Vladimir Pelicic

**Affiliations:** Nanomachines Bactériennes de Virulence (NanoBacVir), Inserm U 1362, Laboratoire de Chimie Bactérienne, CNRS–Aix-Marseille Université UMR 7283, Marseille, France; Université de Paris Cité, INSERM U1151, CNRS UMR8253, Institut Necker-Enfants Malades (INEM), Paris, France; Architecture et Fonction des Macromolécules Biologiques (AFMB), CNRS–Aix-Marseille Université UMR 7257, Marseille, France; Laboratoire Jean Perrin, CNRS–Sorbonne Université UMR 8237, Paris, France; Laboratoire d’Ingénierie des Systèmes Macromoléculaires (LISM), Aix-Marseille Université, CNRS, UMR 7255, Marseille, France

## Abstract

Type 4a pili (T4aP), the most widespread and functionally versatile subtype of the type 4 filaments (T4F) superfamily, are functionalised at their tip by the adhesin PilC/PilY1. Although AlphaFold models predict how this unusually large non-pilin protein is displayed at the pilus tip – by interacting through a short C-terminal stretch of residues with the PilHIJK complex of minor pilins – how PilC/PilY1 drives pilus biogenesis at a stage occurring after filament assembly remains unknown. Here, we answer this question in *Neisseria meningitidis*. After showing that the last 12 residues of PilC, which we name “plug”, are both necessary and sufficient for pilus biogenesis, we demonstrate that the plug interacts with PilK through β-strand augmentation by determining a crystal structure of a PilK-PLUG fusion protein. We establish that this interaction is strong and sequence-specific (by performing binding assays with purified PilK and synthetic peptides) and takes place at a late stage of pilus biogenesis occurring after filament assembly (by showing that extracellular addition of plug peptide to cultures of a *ΔpilC* mutant is able to restore pilus biogenesis by acting outside the cells). Finally, we demonstrate that the plug motif markedly stabilises PilK, providing an explanation for the requirement of PilC in pilus biogenesis and for the ability of a PilK-PLUG fusion protein to bypass this requirement. Together, these findings delineate the mechanism and chronology of events underlying PilC/PilY1 role in pilus biogenesis, reconcile all previous observations about this protein, and establish a broadly applicable model for T4aP biogenesis.

**SIGNIFICANCE STATEMENT:** Type 4a pili (T4aP) are widespread and functionally versatile bacterial surface appendages, functionalised by the tip-located adhesin PilC/PilY1. While PilC/PilY1 is known to drive pilus biogenesis by acting after filament assembly, the underlying mechanism has remained elusive. We demonstrate in *Neisseria meningitidis*, that a C-terminal 12-residue motif within PilC/PilY1, designated the “plug,” is necessary and sufficient for pilus biogenesis. Crystal structure, binding and stability assays reveal that the plug interacts with the minor pilin PilK via β-strand augmentation, sequence-specifically, after filament assembly, and markedly stabilises it. This study delineates the mechanism and chronology of events underlying PilC/PilY1 role in T4aP biogenesis, establishing a broadly applicable model consistent with all previous observations.

## INTRODUCTION

Type 4 filaments (T4F) – a superfamily of nanomachines ubiquitous in Bacteria and Archaea – are helical assemblies of type 4 pilins, assembled by conserved multiprotein machineries (1, 2). Among them, type 4a pili (T4aP) are the best-characterised and have been studied for decades because they are virulence factors in many human bacterial pathogens (3). T4aP are also arguably the most widespread, functionally versatile, and complex T4F. They are found across most phyla of both diderm and monoderm bacteria (2), where they mediate many of the hallmark functions associated with T4F (1), including adhesion, biofilm formation, twitching motility, DNA uptake, and mechanosensing. T4aP biogenesis in diderms cryo-electron tomography (cryo-ET) (4).

One of these conserved components, PilC/PilY1, was first discovered and characterised in the pathogenic *Neisseria* species (*N. gonorrhoeae* and *N. meningitidis*) where it is known as PilC (5). It was subsequently found to be conserved across diverse species, in which it is usually termed PilY1 (6). We thus decided to adopt a unified PilC/PilY1 nomenclature throughout this manuscript. PilC/PilY1 is the largest protein required for T4aP biogenesis, with a molecular weight exceeding 100 kDa. It consists of a variable N-terminus and a conserved C-terminus (7). It is encoded with a signal peptide I (SPI), indicating that it is translocated across the cytoplasmic membrane by the Sec machinery, but PilC/PilY1 consistently co-purifies with pili (5, 6). Immunogold electron microscopy localised PilC/PilY1 at the pilus tip (8, 9), a finding later confirmed by cryo-ET studies (10, 11), which further revealed that PilC/PilY1 interacts with a complex of four minor pilins, conserved across several T4F systems (12). Although these minor pilins are often designated by different names in different species (12), for simplicity, we will refer to the complex as PilHIJK in this manuscript. This complex caps the shaft, which is composed of thousands of copies of the major pilin, and therefore primes filament assembly (13, 14) that proceeds from the tip to the base.

In all species in which it has been characterised, PilC/PilY1 fulfils two distinct functions: it acts both as an adhesin and as a factor required for pilus biogenesis (7). Multiple lines of evidence indicate that these two functions reside in different regions of the protein. The variable N-terminus plays a direct role in adhesion. In *N. meningitidis*, which expresses two paralogs PilC1 and PilC2 that differ mainly by their N-termini, a *ΔpilC1* mutant is severely impaired in adhesion to human endothelial cells, whereas a *ΔpilC2* mutant is unaffected (15). Similarly, in *Kingella kingae*, which also expresses PilC1 and PilC2 paralogs, adhesion assays using purified N-terminal domains showed that PilC1, but not PilC2, mediates adherence to selected extracellular matrix proteins (16). Consistent with these observations, a comprehensive bioinformatic analysis of PilC/PilY1 architectures revealed that the variable N-termini frequently correspond to domains found in a variety of adhesins recognising a wide range of ligands (17). By contrast, how PilC/PilY1 controls pilus biogenesis is not understood, although available evidence implicate the conserved C-terminus. In *N. meningitidis*, where PilC1 and PilC2 possess virtually identical C-terminal regions, piliation is abolished only in a *ΔpilC1ΔpilC2* mutant, whereas each single mutant remains normally piliated (15). Likewise in *K. kingae*, mutants lacking the N-terminal adhesion domains of PilC1 and PilC2 are still piliated, demonstrating that the conserved C-terminus of PilC/PilY1 is sufficient to support pilus biogenesis (16). Critically, studies in multiple species have shown that PilC/PilY1 is dispensable for filament assembly *per se* because piliation can be restored in the absence of PilC/PilY1 when pilus retraction is prevented by a mutation of *pilT*, which encodes the T4aP PilT retraction ATPase (16, 18–20). Therefore, PilC/PilY1 drives pilus biogenesis at a late stage occurring after filament assembly.

Despite the availability of a crystal structure of the *Pseudomonas aeruginosa* PilY1 conserved C-terminus (21) and AlphaFold models of the pilus tip in *Myxococcus xanthus* (22), the mechanism by which PilC/PilY1 drives pilus biogenesis remains mysterious. Indeed, the crystal structure revealed an uninformative β-propeller fold containing a Ca^2+^-binding site (21). Disruption of Ca^2+^ coordination through mutation of chelating residues has no effect on piliation in most species, except in *P. aeruginosa*, indicating that this site is not widely essential for function (21, 23, 24). On the other hand, the AlphaFold models in *M. xanthus* that predicted how PilC/PilY1 interacts with the PilHIJK complex (22) do not explain why PilC/PilY1 is required for pilus biogenesis. Here, we address this outstanding question in *N. meningitidis* one of the principal T4aP model systems, using a multidisciplinary approach combining structural modelling, molecular genetics, protein biochemistry and structural biology. Our work establishes a broadly applicable model that delineates the mechanism underlying PilC/PilY1 role at a stage in pilus biogenesis and reconciles previous observations on PilC/PilY1 across diverse T4aP systems.

## RESULTS

### A short C-terminal plug motif in PilC/PilY1 anchors this protein to the tip-localised PilHIJK complex

Our parental strain of *N. meningitidis*, derived from the serogroup C clinical isolate 8013 (25), encodes two PilC/PilY1 paralogs (Fig. 1A), PilC1 and PilC2 (Table S1). Bioinformatic analyses confirm that both proteins start with a SPI and have mature molecular masses of 108 and 110 kDa, respectively. Their N-termini share only 54% sequence identity, whereas their C-termini are 93% identical and encompass the IPR008707 β-propeller domain (Fig. 1A), a signature domain invariably found in PilC/PilY1 proteins (7). By contrast, no known protein domain could be identified in the N-terminal region of PilC1 and PilC2.

**Fig. 1.**
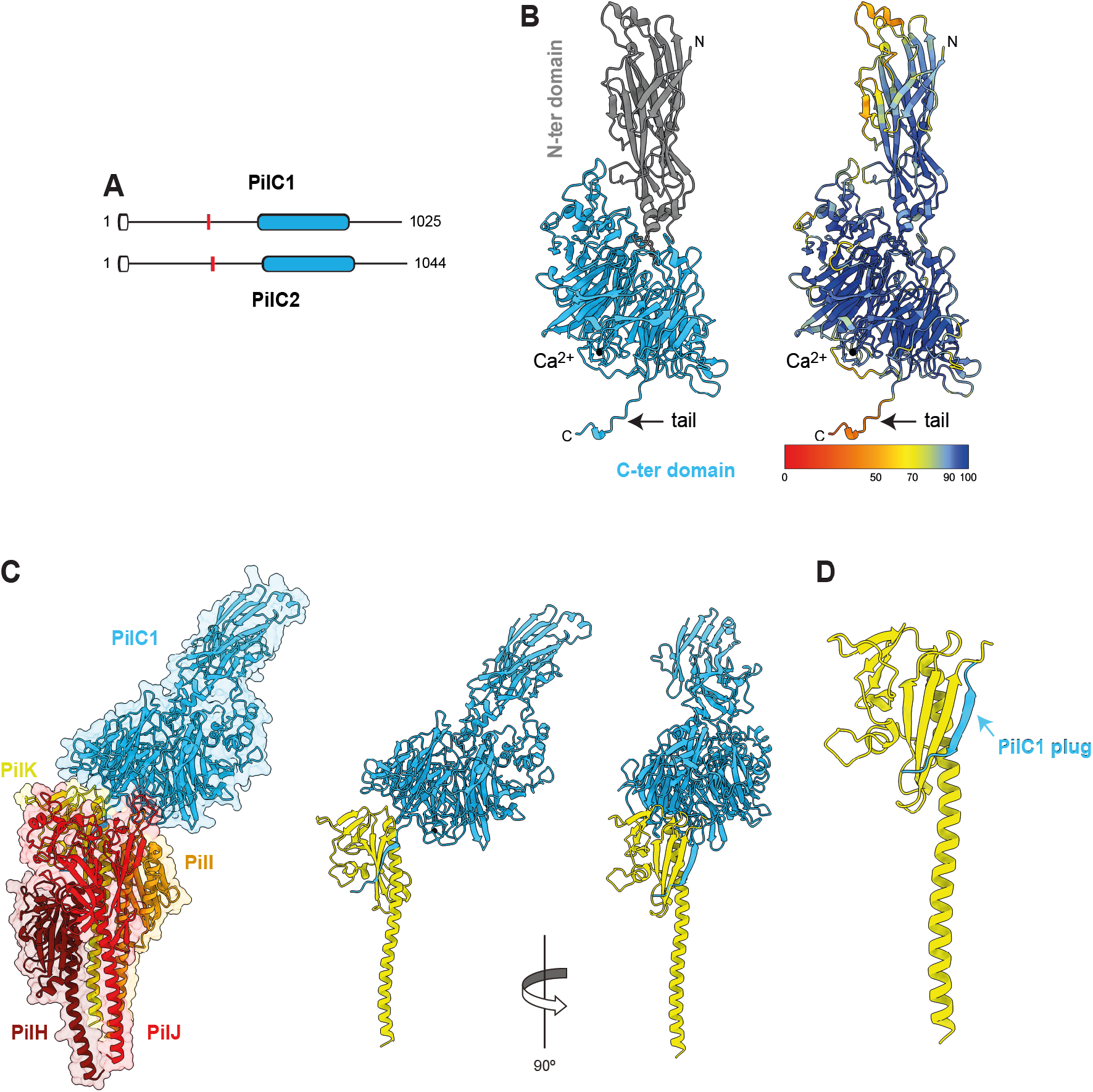
AlphaFold models predict that PilC1/2 of *N. meningitidis* localises to the tip of T4aP via a short C-terminal plug motif that interacts with the PilHIJK complex of minor pilins through β-strand addition to PilK. **A)** Predicted architectures – drawn to scale – of the PilC1 and PilC2 paralogs in *N. meningitidis* strain 8013. This cartoon highlights the typical features of PilC/PilY1 proteins (7): a very large size, a periplasmic localisation via a SPI (white rounded rectangle), a variable N-terminal region (the vertical red line indicates the boundary between N-and C-terminal regions), and a conserved C-terminal region centred on an IPR008707 β-propeller domain (blue rounded rectangle). Panels B, C, D) show the AlphaFold 3 (26) predictions for PilC1 with a bound Ca^2+^ ion as observed in the crystal structure of *P. aeruginosa* PilY1 (21). Because PilC2 models are highly similar, they are shown in a separate Fig. S1 for a sake of clarity. **B**) PilC1 model. Left panel, PilC1 is an elongated protein with two distinct domains. The conserved C-terminal domain includes a bound Ca^2+^ ion and ends with a protruding 12-residue tail. Right panel, this tail is intrinsically disordered as suggested by its very low pLDDT confidence scores. The pLDDT colour key is provided at the bottom. **C**) PilC1HIJK model. PilC1 binds to the tip-localised PilHIJK complex of conserved minor pilins through its 12-residue tail that adopts a β-strand conformation, forming a plug that docks into PilK through β-strand augmentation (28). **D**) Zoom on the interaction between PilK and the plug motif of PilC1. The plug extends the second β-sheet in the globular head of PilK through an extensive network of hydrogen bonds and salt bridges with its terminal β-strand.

To gain structural insight into PilC1/2, we used AlphaFold 3 (26) to predict the structures of mature proteins with a bound Ca^2+^ ion, as observed in the crystal structure of the β-propeller domain of *P. aeruginosa* PilY1 (21). Both PilC1 (Fig. 1B) and PiC2 (Fig. S1A) models were generated with high confidence (pTM scores of 0.9 and 0.91 for PilC1 and PilC2, respectively) and are remarkably similar (Fig. S1B). Overall, PilC1/2 are elongated proteins composed of two distinct domains, the C-terminal of which binds the Ca^2+^ ion and ends in a 12-residue “tail”. This tail, identical in sequence in both proteins, is the region with the lowest pLDDT confidence score in PilC1 (Fig. 1B) and PiC2 (Fig. S1A), suggesting that it is intrinsically disordered. Searches for structural homologs of PilC1/2 in the Protein Data Bank (PDB) unsurprisingly identified the C-terminal β-propeller domain of *P. aeruginosa* PilY1 (21) (PDB 3XH6) as the closest structural match of the C-terminal regions of PilC1/2 (Fig. S1C). The major difference is that PilC1/2 has an extra blade compared to PilY1 that was described as a seven-bladed β-propeller (21). The Ca^2+^-binding site, that is situated in a loop between two β-strands of blade 4, is extremely conserved (Fig. S1D). As for PilC1/2 N-terminal regions, the best structural match is the fimbrial adhesin AtfE from *Proteus mirabilis* (PDB 6H1Q), a predicted lectin (27), which suggests that PilC1/2 bind carbohydrates (Fig. S1E).

Next, we predicted the structures of complexes between PilC1 or PilC2, each with a bound Ca^2+^ ion, and the four minor pilins PilHIJK. Both PilC1HIJK (Fig. 1C) and PilC2HIJK (Fig. S2A) complexes were predicted with high confidence, with ipTM scores of 0.84 and 0.81, respectively. The two models, which are highly similar and consistent with the previously reported models of the *M. xanthus* pilus tip (22), predict that the tail residues of PilC1/2 behave as a “plug” anchoring this protein to the PilHIJK complex (Fig. 1C and Fig. S2A). Unlike when PilC1/2 are modelled alone, the plug motif adopts a β-strand conformation and extends the β-sheet in the globular head of PilK (Fig. 1D). Although this mode of protein-protein interaction, known as β-strand augmentation (28), is unusual in T4F, it is conserved across T4aP for PilC/PilY1 (Fig. S2B). To identify the interacting residues in *N. meningitidis* PilC1HIJK and PilC2HIJK models and define the types of interaction they establish, we used several bioinformatic tools. These analyses showed that PilC1/2 interact almost exclusively with PilK – and to a lesser extent with PilJ – through an extensive network of hydrogen bonds and salt bridges (Supplementary dataset 1). An overwhelming majority of these contacts are mediated by residues within the PilC1/2 plug motif.

These and previously published modelling data (22) predict that a short C-terminal plug motif in PilC/PilY1, representing merely ∼1% of the protein, is the principal determinant of interaction with the tip-localised PilHIJK complex.

### The PilC/PilY1 plug motif is both necessary and sufficient for pilus biogenesis

The above modelling data predict that the PilC/PilY1 plug motif is likely to be functionally important. To test this hypothesis in *N. meningitidis*, we first generated a strain suitable for functional analysis. Since meningococci encode two paralogs, PilC1 and PilC2, that are functionally redundant in pilus biogenesis (15) and whose expression is subject to frequent phase variation (5), we constructed a strain carrying markerless deletions of both *pilC1* and *pilC2*. As expected, the *ΔpilC1ΔpilC2* mutant was non-piliated (Fig. 2A) and non-competent for genetic transformation (Fig. 2B). Piliation and transformation were restored by complementation with a wild-type (WT) *pilC1* allele not subject to phase variation, expressed from an anhydrotetracycline (aTC)-inducible P*tet* promoter and integrated ectopically in the chromosome (29) (Fig. 2A and Fig. 2B). In contrast, complementation with a mutant allele leading to the synthesis of a PilC1_Δplug_ variant lacking the 12-residue plug motif failed to restore either piliation or transformation in the *ΔpilC1ΔpilC2* mutant (Fig. 2A and Fig. 2B). These results demonstrate that the C-terminal plug motif in PilC1/2 is necessary for pilus biogenesis.

**Fig. 2.**
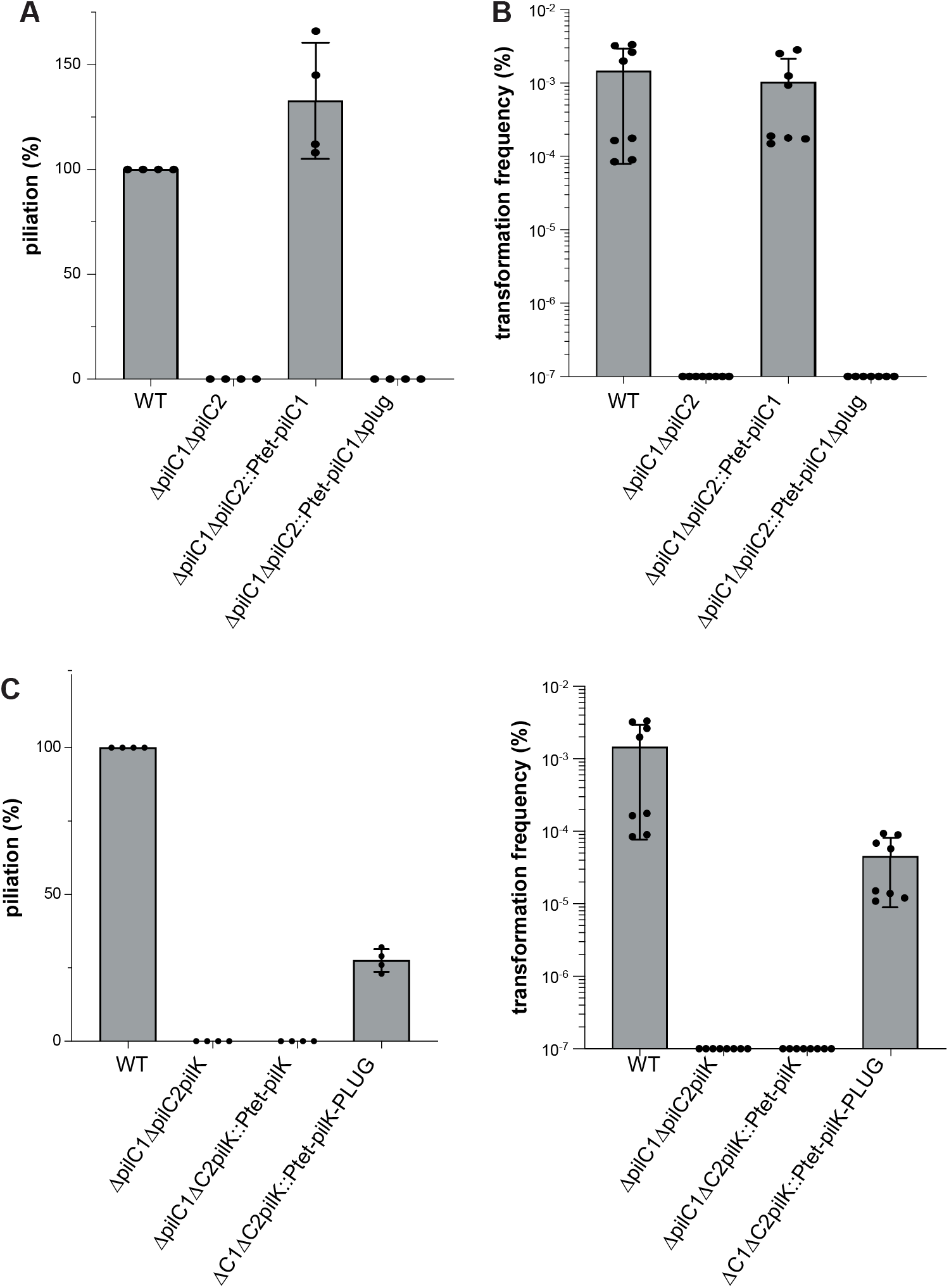
The C-terminal plug motif of PilC1/2 is both necessary and sufficient for the biogenesis of functional T4aP in *N. meningitidis*. Because *N. meningitidis* strain 8013 encodes paralogous *pilC1* and *pilC2* genes whose expression is subject to phase variation (5) and that are functionally redundant in pilus biogenesis (15), we first constructed a markerless *ΔpilC1ΔpilC2* mutant. This mutant was complemented with diverse *pilC1* alleles not subject to phase variation, expressed from an aTC-inducible P*tet* promoter and ectopically integrated in the chromosome (29). Specifically, the different alleles encoded either PilC1 or a PilC1_Δplug_ derivative lacking the 12-residue plug. **A**) Pilus biogenesis was assayed by quantifying piliation by immunofluorescence using a monoclonal antibody specific for the filaments of our parental strain of *N. meningitidis* (60). Results are expressed in % piliation relative to WT, which is set to 100%, and are the average ± standard deviation from four independent experiments. **B**) Pilus functionality was assayed by quantifying transformation frequencies using genomic DNA carrying a cassette promoting resistance to nalidixic acid. Results are expressed as % of recipient cells transformed and are the average ± standard deviation from eight independent experiments. **C**) Expression of a PilK-PLUG fusion protein in *N. meningitidis* bypasses the requirement for PilC1/2 in pilus biogenesis. We constructed a *ΔpilC1ΔpilC2pilK* triple mutant and complemented it with *pilK* or *pilK-PLUG* alleles expressed from an aTC-inducible P*tet* promoter and ectopically integrated in the chromosome (29). Piliation levels (left panel) and frequency of transformation (right panel) were measured as above. Values expressed in % correspond to mean ± standard deviation from four and eight independent experiments for piliation and transformation, respectively

Next, since the plug motif in PilC1/2 is the principal determinant of interaction with the tip-localised PilK (Supplementary dataset 1), we tested whether expression of a chimeric PilK-PLUG fusion protein in *N. meningitidis*, in which the plug motif of PilC1/PilC2 is fused to the C-ter of PilK, might be sufficient for pilus biogenesis in the absence of PilC1/2. We constructed a *ΔpilC1ΔpilC2pilK* triple mutant and complemented it with aTC-inducible *pilK* or *pilK-PLUG* alleles, ectopically integrated in the chromosome (29). Complementation with *pilK-PLUG* restored significant but not WT levels of piliation and natural transformation (Fig. 2C), whereas complementation with *pilK* failed to restore either phenotype. This demonstrates that expression of a PilK-PLUG fusion protein in *N. meningitidis* bypasses, at least in part, the requirement for PilC1/2 in pilus biogenesis.

Taken together, these molecular genetic data show that the PilC/PilY1 plug is both necessary and sufficient for pilus biogenesis.

### The PilC/PilY1 plug interacts specifically with PilK through β-strand augmentation

Although all AlphaFold models predict that the PilC/PilY1 plug interacts with PilK through β-strand augmentation, this remains to be structurally demonstrated. To achieve this, we generated and purified a chimeric 6His-PilK-PLUG in *Escherichia coli* and determined its 3D structure by X-ray crystallography. This protein lacks the first 24 hydrophobic residues of mature PilK from *N. meningitidis*, which were replaced by a 6His tag. This strategy is commonly used to improve pilin solubility without altering the protein structure (30). The 6His-PilK-PLUG protein crystallized readily and, following optimisation of crystal growth conditions, its structure was solved at 2 Å resolution by molecular replacement (Fig. 3). The asymmetric unit contains six monomers arranged as two trimeric assemblies (Fig. S3). The structure revealed a characteristic pilin fold (31), consisting of a long N-terminal α1-helix packed against a globular head built around a β-sheet scaffold composed of a first three-stranded antiparallel β-sheet (interspersed with extended loops), followed by a long loop containing a 3_10_ helix, and ending in a second four-stranded antiparallel β-sheet (Fig. 3A). Critically, the second β-sheet of 6His-PilK-PLUG is extended by the antiparallel incorporation of the plug motif of PilC/PilY1, which adopts a β-strand conformation as predicted. Superposition of the crystal structure with the corresponding portion of PilC1HIJK model is excellent (RMSD = 0.629 Å) (Fig. 3B), which validates the mode of interaction between PilC/PilY1 and PilK that was predicted by AlphaFold.

**Fig. 3.**
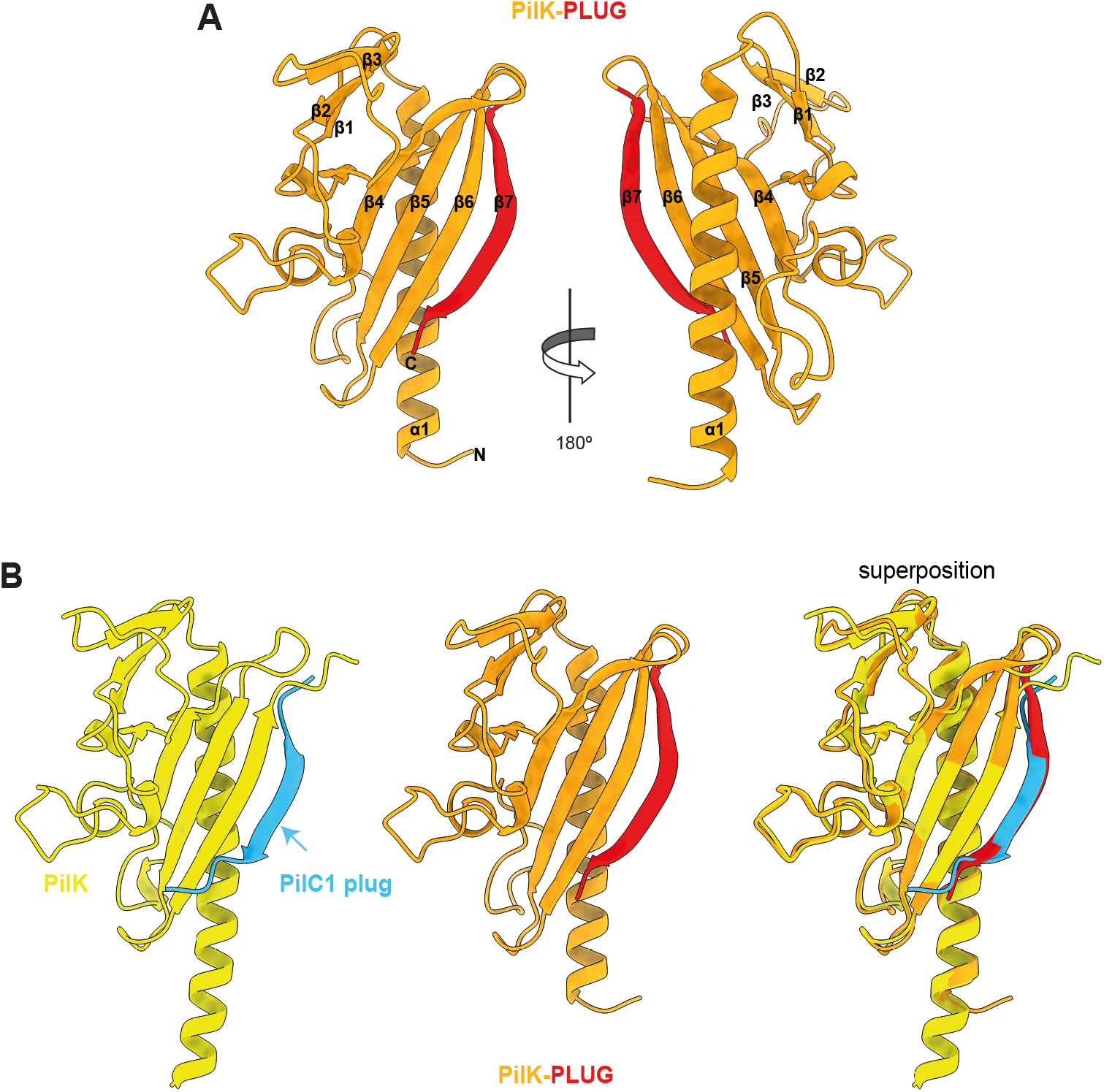
The crystal structure of a PilK-PLUG fusion protein demonstrates that the interaction between the plug motif of PilC/PilY1 and PilK is mediated through β-strand augmentation. **A)** Crystal structure of PilK-PLUG. The structure adopts a characteristic pilin fold with of an N-terminal α-helix packed against a globular head built around two antiparallel β-sheets. The first β-sheet is three-stranded, whereas the second is four-stranded. The last strand (β7) of this second β-sheet is formed by the fused PLUG that is highlighted in red. Left and right panels show orthogonal views. **B)** Comparison of the crystal structure with the PilHIJKC1 model. Only the portion of the model corresponding to the PilK-PLUG construct is shown and the same colour code than in Fig. 1C is used. The excellent superposition validates the mode of interaction predicted by AlphaFold between PilK and the plug motif of PilC/PilY1.

To establish the strength and specificity of the plug binding to PilK, we used bio-layer interferometry (BLI), a label-free optical analytical technique that measures real-time biomolecular interactions (32). We produced 6His-PilK in *E. coli* and investigated its interaction with various plug-derived synthetic 12-residue peptides (Table S2). The PLUG peptide corresponds to the native plug sequence, GLUP contains the same residues in a different order, whereas PLUG+/-carries a permutation of two oppositely charged residues, predicted to form intermolecular salt bridges with PilK (Supplementary dataset 1). After immobilising purified 6His-PilK on the biosensor, we first monitored the association and dissociation kinetics of the PLUG peptide at increasing concentrations. Analysis of these binding curves yielded an equilibrium dissociation constant (*K*d) of 61.95 ± 31.4 µM for the interaction between PilK and the PLUG peptide (Fig. 4A). We next compared the binding of all three peptides at a concentration of 50 µM, close to the *K*_d_ determined for PLUG. PilK displayed an affinity for the PLUG peptide consistent with the experiment above, which was markedly higher than for either GLUP or PLUG+/-(Fig. 4B). No interaction was detected with PLUG+/-, whereas for the interaction with GLUP the signal was very weak, close to the limit of detection of the instrument. Together, these results demonstrate that the C-terminal plug motif of PilC/PilY1 binds PilK strongly and specifically.

**Fig. 4.**
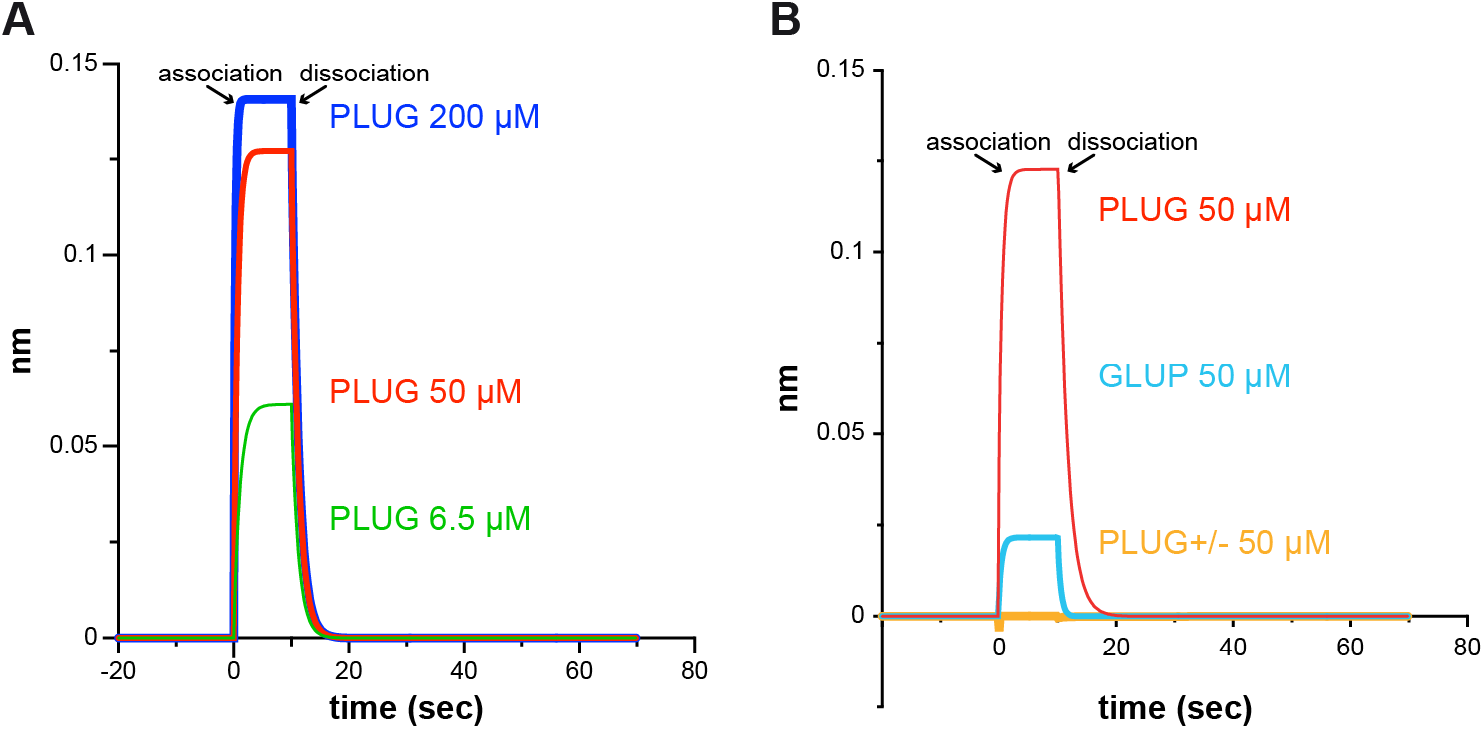
PilK interacts with the C-terminal plug motif of PilC/PilY1 in a strong and specific fashion. BLI was used to investigate the interaction of purified 6His-PilK with plug-derived 12-residue synthetic peptides. The purified protein was immobilised on biosensors, which were then immersed in wells containing the peptides. The PLUG peptide corresponds to the native plug sequence, GLUP contains the same residues in a different order, whereas PLUG+/-carries a permutation of two oppositely charged residues, predicted to form intermolecular salt bridges with PilK (Supplementary dataset 1). **A**) Binding curves of increasing concentrations of PLUG peptide (6.5, 50 and 200 µM) to 6His-PilK. **B**) Comparative binding curves of either PLUG, GLUP, or PLUG+/-peptides (all at 50 µM) to 6His-PilK.

Taken together, these structural and biochemical data demonstrate that the PilC/PilY1 plug interacts specifically with PilK through β-strand augmentation.

### The PilC/PilY1 plug motif acts after pilus assembly

Because PilC/PilY1 drives pilus biogenesis at a stage occurring after filament assembly (16, 18–20), we next determined when the plug motif was involved in pilus biogenesis. To do this we asked whether piliation and transformation could be restored in the *ΔpilC1ΔpilC2* mutant upon extracellular addition of the above synthetic peptides. Remarkably, addition of the PLUG peptide to cultures of the *ΔpilC1ΔpilC2* mutant restored substantial levels of both piliation (Fig. 5A) and natural transformation (Fig. 5B). This complementation was dependent on the presence of PilK because the PLUG peptide failed to restore either phenotype in a *ΔpilC1ΔpilC2pilK* mutant (Fig. 5A and Fig. 5B). The complementation was highly specific of the PLUG sequence since GLUP and PLUG+/-peptides failed to restore either phenotype (Fig. 5A and Fig. 5B). Together, these findings show that the extracellular addition of a synthetic PLUG peptide is sufficient for pilus biogenesis in a *ΔpilC1ΔpilC2* mutant and that this activity depends both on the plug native sequence and the presence of PilK.

**Fig. 5.**
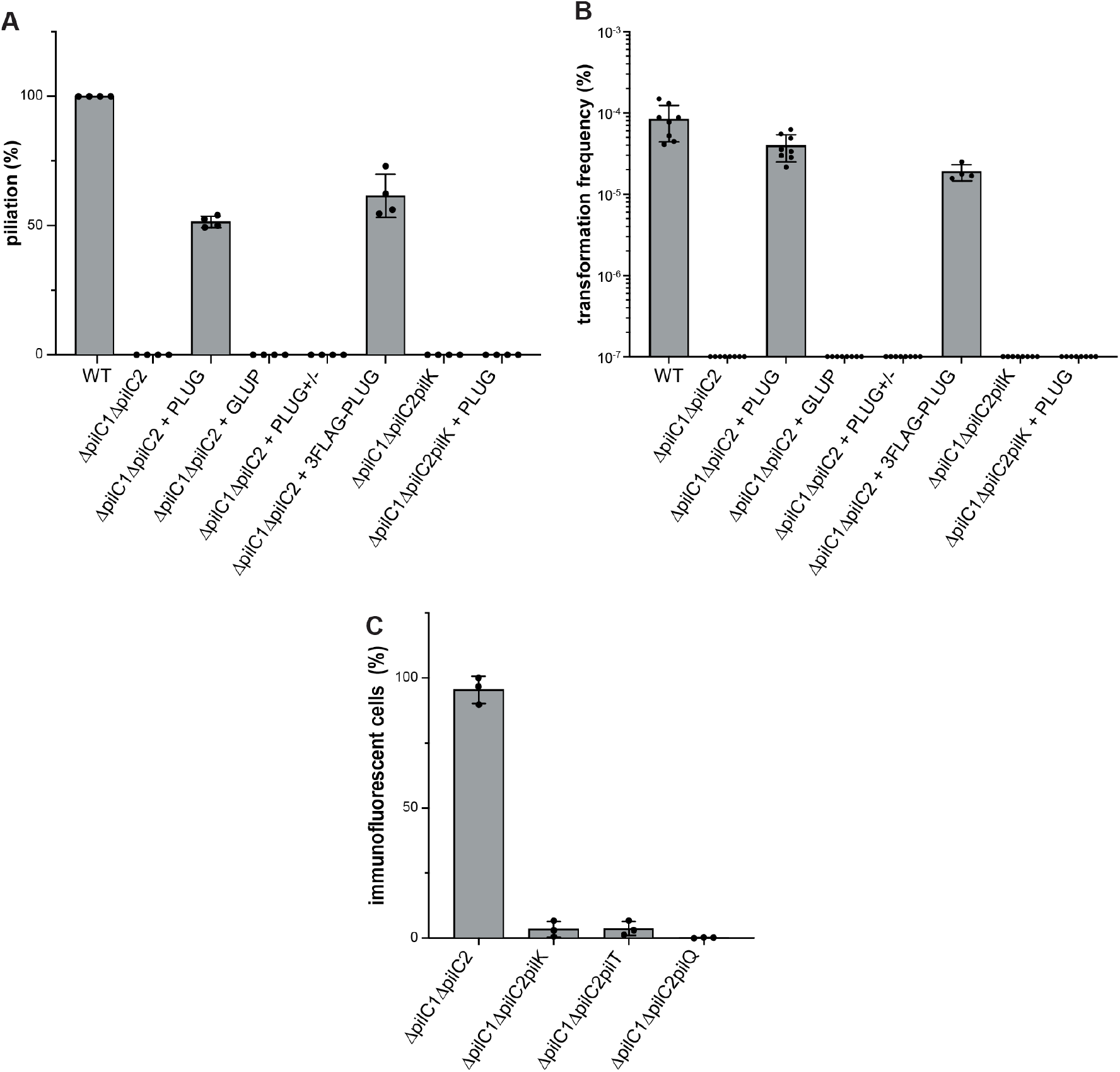
Extracellular addition of plug peptide restores the biogenesis of functional T4aP in a *ΔpilC1ΔpilC2* mutant by acting after pilus assembly. To determine at which stage of pilus biogenesis the plug motif was involved, we asked whether piliation and transformation could be restored in various *ΔpilC1ΔpilC2* mutants upon extracellular addition of synthetic PLUG, GLUP, or PLUG+/-peptides. We also tested 3FLAG-PLUG, a PLUG peptide with three FLAG tags at its N-terminus. **A)** Pilus biogenesis was assayed by quantifying piliation as above. Results are expressed in % piliation relative to WT, which is set to 100%, and are the average ± standard deviation from four independent experiments. **B)** Pilus functionality was assayed by quantifying transformation frequencies as above. Results are expressed as % of recipient cells transformed and are the average ± standard deviation from eight independent experiments (four for 3FLAG-PLUG). **C**) Peptide uptake assays. Intracellular 3FLAG-PLUG peptide was detected by immunofluorescence using a specific anti-FLAG antibody, upon bacterial permeabilization. We tested the WT, and three triple mutants *ΔpilC1ΔpilC2pilK*, *ΔpilC1ΔpilC2pilT* and *ΔpilC1ΔpilC2pilQ*. Results are expressed as % of immunofluorescent cells and are the average ± standard deviation from three independent experiments.

Next, we tracked the synthetic PLUG peptide to determine if it acts outside or inside the cells. To do this, we synthesized a 3FLAG-PLUG peptide, which restored piliation and natural transformation as efficiently as the unmodified PLUG peptide (Fig. 5A and Fig. 5B). Immunofluorescence analysis of permeabilised bacteria using a specific anti-FLAG antibody showed that nearly all *ΔpilC1ΔpilC2* cells were fluorescent and thus internalised the 3FLAG-PLUG peptide, whereas very few *ΔpilC1ΔpilC2pilK* cells were fluorescent (Fig. 5C). By contrast, no intracellular fluorescence was detected in a *ΔpilC1ΔpilC2pilQ* mutant (Fig. 5C), in which surface-exposed pili are not observed because of the lack of secretin PilQ, or a *ΔpilC1ΔpilC2pilT* mutant (Fig. 5C), which cannot retract its pili because it lacks the PilT retraction ATPase (33). These observations strongly suggest that the synthetic PLUG peptide acts by engaging the pilus tip from the extracellular milieu.

Taken together, these data demonstrate that the PilC/PilY1 plug functions at a late stage in pilus biogenesis occurring after filament assembly.

### The PilC/PilY1 plug motif drives pilus biogenesis by stabilising PilK

While the previous results demonstrate the mode and chronology of PilC/PilY1 interaction with PilK, they do not explain why PilC/PilY1 is required pilus biogenesis. We reasoned, based on two well-known facts, that this might be due to a stabilising effect of the PilC/PilY1 plug on PilK. First, two interacting proteins frequently stabilise one another. Second, since PilK is required for piliation, its impaired stability in the absence of PilC/PilY1 would have a negative impact on pilus biogenesis. To test whether the plug motif stabilises PilK, we first compared the thermal stability of purified 6His-PilK and 6His-PilK-PLUG proteins by performing Thermofluor assays (34). This assay monitors protein unfolding by measuring the fluorescence of a hydrophobic dye that binds to the hydrophobic protein core upon denaturation. The melting curves revealed a striking increase in the thermal stability of PilK-PLUG relative to PilK, with melting temperatures (Tm) of 58.37 ± 0.23°C and 45.59 ± 0.21°C, respectively (Fig. 6A).

**Fig. 6.**
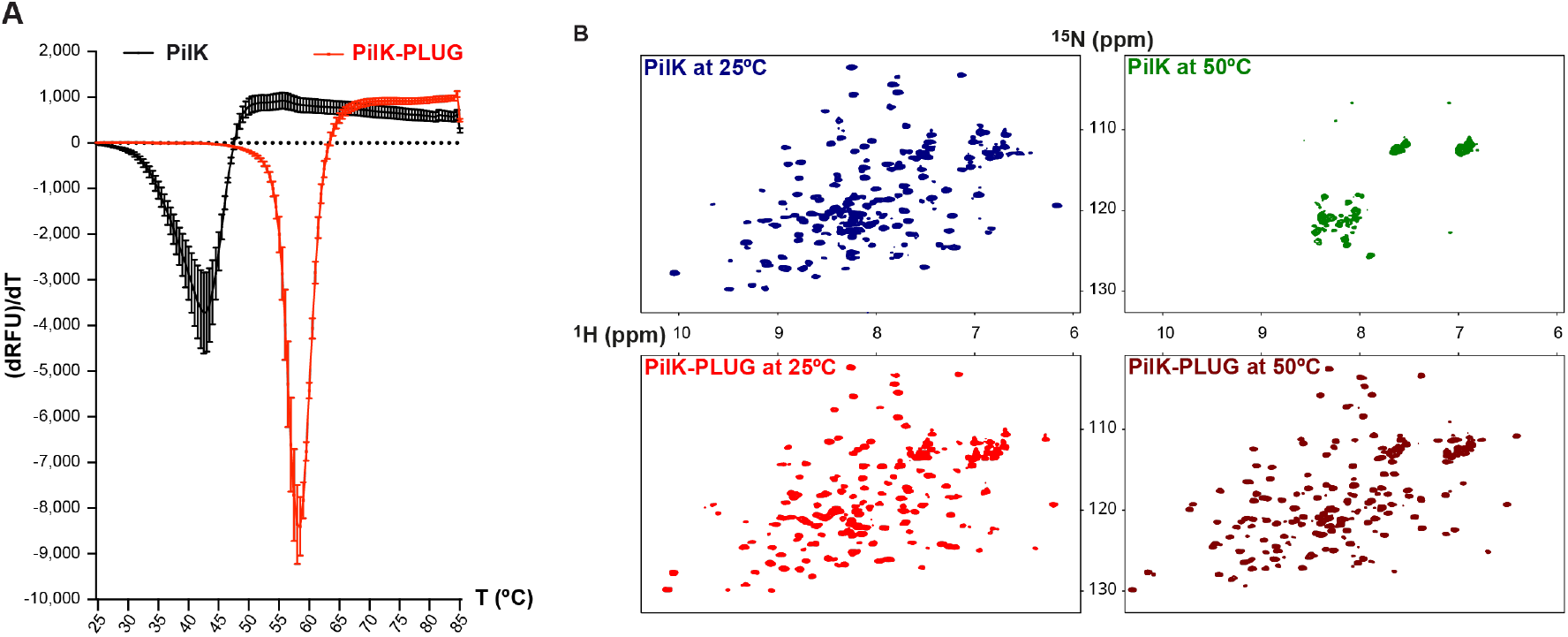
The PilC/PilY1 plug motif strongly stabilises PilK. **A**) Assessing thermal stability for 6His-PilK and 6His-PilK-PLUG using Thermofluor (34). The purified proteins were heated in the presence of a hydrophobic dye SYPRO Orange. Melting curves are represented using the first derivative (dRFU)/dT of the raw fluorescence data. The Tm corresponds to the apex. The values are the average ± standard deviation from three independent experiments. **B**) Assessing 6His-PilK and 6His-PilK-PLUG structural integrity and stability at different temperatures by NMR. The ^1^H-^15^N HSQC spectra for PilK (upper panels) and PilK-PLUG (lower panels) were recorded at either 25°C (left panels) or 50°C (right panels).

Next, we assessed the structural integrity and stability of 6His-PilK and 6His-PilK-PLUG directly by nuclear magnetic resonance (NMR), by recording ^1^H-^15^N Heteronuclear Single Quantum Coherence (HSQC) spectra at 25°C and 50°C (Fig. 6B). Analysis of 2D NMR spectra provides a characteristic fingerprint and enables a direct assessment of protein quality, folding status and dynamics. At 25°C, both proteins displayed well-dispersed NMR signals, consistent with folded conformations, whereas the spectra diverged markedly at 50°C. At this higher temperature, the PilK spectrum collapsed into a narrow distribution of resonances characteristic of an unfolded protein, consistent with the Tm measured by Thermofluor. By contrast, the PilK-PLUG spectrum remained well dispersed, indicating that this protein retained its folded conformation (Fig. 6B). We also compared the stability of the two purified proteins at 25°C by monitoring the HSQC spectra over time (Fig. S4). In contrast to the PilK-PLUG spectrum that showed remarkable preservation over 33 h, indicative of high protein stability, we observed noticeable changes in peak intensities and positions for PilK during the same time, indicating that this protein undergoes significant structural and/or dynamic alterations (Fig. S4). This was particularly noticeable in the central region of the PilK spectrum, which therefore contains resonances from highly dynamic residues. Signals in this region exhibited a marked increase in intensity over time, consistent with an increase of highly flexible and/or unfolded conformations in the population. In addition, selected resonances outside the central region that correspond to well-folded segments of the protein, showed a decrease in intensity over time, consistent with a progressive decrease in sample stability (Fig. S4).

Together, these data demonstrate that PilC/PilY1 drives pilus biogenesis by stabilising PilK, which is an essential component of the PilHIJK minor pilin complex that primes pilus assembly (13, 14).

## DISCUSSION

T4aP are remarkably versatile filaments found across most phyla of both diderm and monoderm bacteria (2), and have been studied for decades because they act as virulence factors in many human bacterial pathogens (3). Although studies of T4aP have provided much of our current understanding of the T4F superfamily to which they belong (1), fundamental aspects of pilus biogenesis or pilus-mediated functions remain poorly understood. Here, by focusing on PilC/PilY1, a conserved component of the complex T4aP biogenesis machinery (35, 36) that acts both as an adhesin and as a factor required for pilus biogenesis (7), we report findings that significantly advance our understanding of T4aP and T4F biology.

First, whereas it has been long known that PilC/PilY1 functionalises T4aP by localising to the pilus tip (8, 9), it is only more recently that cryo-ET studies revealed that this unusually large non-pilin protein interacts with the tip-localised PilHIJK complex (10, 11). AlphaFold models in *M. xanthus* (22) predicted that a short stretch of C-terminal residues in PilC/PilY1, which we termed the plug, recruits this protein to the pilus tip by interacting with PilK. Here, using AlphaFold 3 models of substantially improved accuracy (26) and molecular genetics in *N. meningitidis*, we demonstrate the critical importance of the plug for pilus biogenesis, showing that is both necessary (a PilC1_Δplug_ variant failed to restore either piliation or transformation in a *ΔpilC1/2* mutant) and sufficient (a PilK-PLUG fusion protein bypasses the requirement for PilC1/2 in pilus biogenesis). Importantly, while this manuscript was under consideration, two studies used similar AlphaFold models as blueprints to report similar results in other species (37, 38). Ellison *et al*. demonstrated both in *N. gonorrhoeae* and *Acinetobacter baylyi* that mutants expressing a PilC/PilY1_Δplug_ variant phenocopy non-piliated *ΔpilC/pilY1* mutants and that expression of PilK-PLUG fusion proteins partly compensate for the absence of PilC/PilY1 (37). Yount *et al*. demonstrated in *K. kingae* that truncation of the plug motif in PilC1/2 or point mutations targeting conserved residues both in the plug and in PilK significantly impair pilus biogenesis and/or pilus-mediated functions (adherence to cells, twitching motility and natural transformation) without altering PilC1/2 protein expression (38). These studies strongly support the cross-species generalisation of the mechanism we propose below.

Second, although all AlphaFold models generated by us and others (22, 37–39) agree that the interaction between PilC/PilY1 and PilK occurs through β-strand augmentation, here we demonstrate for the first time that these predictions are correct. Our crystal structure of *N. meningitidis* PilK-PLUG fusion protein shows that the PilC1/2 plug indeed adopts a β-strand conformation and extends the second β-sheet of the PilK globular head through β-strand augmentation (28). Despite the structural conservation in different models, the plug motifs in different systems show little apparent sequence conservation. It is likely that the interacting residues within the PilC/PilY1 plug and the C-terminal β-strand of PilK have co-evolved in each system to preserve both the strength and specificity of the interaction. This is consistent with our BLI data demonstrating that binding of PLUG peptides to PilK is strong and sequence-specific. This mechanism of protein presentation at the T4aP tip further expands the diversity of strategies employed by T4F to functionalise this critical region of the filament. Indeed, previously described mechanisms differ substantially. In monoderm T4aP, which lack PilC/PilY1 and only have a structural homologue of PilI, the tip is functionalised by large modular minor pilins bearing diverse C-terminal modules that bind a variety of ligands (17, 40). Interestingly, many of the C-terminal domains in these modular pilins are also found at the N-terminus in PilC/PilY1 (17). This observation suggests that monoderm and diderm T4aP have converged to use the same evolutionary tinkering strategy – recruitment of diverse ligand-binding domains to the pilus tips – but employ distinct tip-presentation modules, either a pilin or an IPR008707-containing domain ending with a plug motif. By contrast, T4bP and T4dP functionalise their tips for purposes unrelated to adhesion. T4bP, which are restricted to diderms and considerably less widespread than T4aP (2), recruit non-pilin “cargo” proteins for secretion through a tip-localised homo-trimer of a modular minor pilin. In this system, the extended N-terminus of the cargo protein acts as an export signal by binding within the cleft between two minor pilins in the homo-trimer (41, 42). In T4dP, which mediate DNA uptake in hundreds of monoderm species (43), the PilHIJK complex itself has evolved to bind extracellular DNA through the interface between PilH and PilJ (44, 45).

Third, our study presents two important and new lines of evidence converging to reveal the molecular mechanism underlying PilC/PilY1 role in pilus biogenesis, which had long been recognised to occur after filament assembly but remained mysterious. Indeed, almost three decades ago it has been reported in *N. gonorrhoeae* that an absolute defect in T4aP biogenesis in a *pilC* mutant can be suppressed by loss-of-function mutations in *pilT* (18), which was later confirmed in multiple other model species (19). The first line of evidence is the demonstration that extracellular addition of a synthetic PLUG peptide to cultures of the *ΔpilC1ΔpilC2* mutant restores piliation. Importantly, this activity, which depends on the presence of PilK, results from a binding event occurring outside the cells and therefore after filament assembly. The second line of evidence is the finding that PilK is dramatically stabilised upon interaction with the PilC/PilY1 plug motif. This allows us to propose a unified model for T4aP biogenesis that reconciles all previous observations about PilC/PilY1 (Fig. 7).

**Fig. 7.**
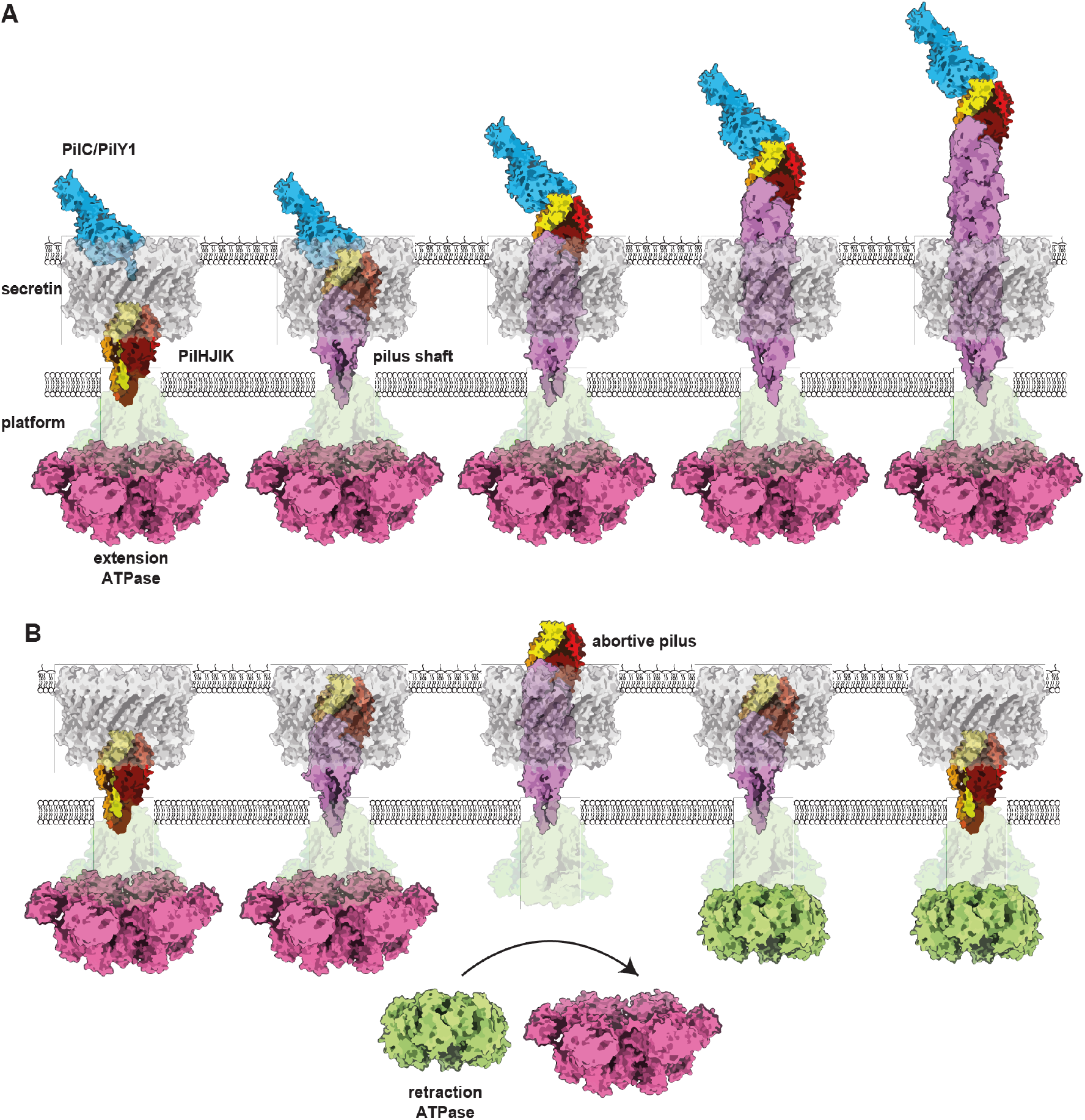
Mechanistic model explaining how PilC/PilY1 drives T4aP biogenesis by acting at a step occurring after filament assembly. Cartoons are based on AlphaFold 3 (26) predictions with *N. meningitidis* strain 8013 proteins (Table S1), except the for the pilus shaft for which we used the cryo-EM structure PDB 5KUA (75). The secretin (PilQ) was modelled as a tetradecamer (76), using only the oligomerisation/membrane-spanning domain (77). The platform protein (PilG) was modelled as a trimer, whereas the extension (PilF) and retraction (PilT) ATPases were modelled as hexamers. **A**) Unimpeded pilus biogenesis in the presence of PilC/PilY1. **B**) Abortive pilus biogenesis in the absence of PilC/PilY1.

In our model, pilus biogenesis begins with assembly of the PilHIJK priming complex (13, 14) within the T4aP machinery at the cytoplasmic membrane, an event that has been visualised by cryo-ET (4, 10, 11). Then, incorporation of major pilin subunits underneath this priming complex drives its upward movement towards the secretin in the outer membrane (Fig. 7A and Fig. 7B). When PilC/PilY1 is present in the periplasm, it is preloaded in the secretin channel, which it seals like a “champagne cork” as visualised by cryo-ET (11). Preloaded PilC/PilY1 is positioned with its C-terminal plug domain facing the growing pilus (Fig. 7A). Once the pilus reaches this preloaded PilC/PilY1, it engages its plug through β-strand augmentation of the PilK subunit at the apex of the PilHIJK complex (Fig. 7A). Because the plug markedly stabilises PilK, it is likely that this in turn stabilises the PilHIJK priming complex and thereby the growing filament, promoting unimpeded pilus extension (Fig. 7A). In the absence of PilC/PilY1, we surmise that the absence of PilK stabilisation leads to a lower stability of the PilHIJK complex, which sends a signal that is somehow sensed by the cells through the filament, shifting the balance from pilus extension towards pilus retraction by promoting exchange of motors at the base (Fig. 7B). Strikingly, even in the absence of PilC/PilY1, filaments can still be exported to the surface (Fig. 7B) as demonstrated here by the ability of a synthetic PLUG peptide to restore pilus biogenesis in a *ΔpilC1ΔpilC2* mutant by engaging the pilus tip from the extracellular milieu. However, because the *ΔpilC1ΔpilC2* mutant is non-piliated its filaments must be too short and/or retracted too rapidly to be detectable, a process we termed abortive pilus biogenesis. This is consistent with the observations that abortive pilus biogenesis in the absence of PilC/PilY1 can be overridden either by abolishing pilus retraction through a loss-of-function mutation in *pilT*, as described in many previous studies (16, 18–20), or by expressing a more stable PilK-PLUG fusion protein as shown in this study.

In conclusion, this study explains how PilC/PilY1, a conserved protein required for T4aP biogenesis at a step occurring after filament assembly (18, 19), drives pilus biogenesis through a short C-terminal plug motif. This elegant mechanism provides new opportunities both for bioengineering applications by exploiting the plug motif to display useful protein cargos at the filament tip, and for therapeutic interventions by interfering with T4aP biogenesis. Given the widespread distribution of T4aP across bacteria (2) and their central role in the virulence of many human pathogens (3), these findings have broad implications for bacterial biology.

## MATERIAL AND METHODS

### AlphaFold and bioinformatics

We used AlphaFold 3 (26) for modelling protein 3D structures. Modelling was done using the mature form of the proteins in which the predicted SPI and SPII were manually removed from the protein sequences downloaded from the UniProtKB (46) (Table S1). We privileged strains in which T4aP have been characterised and/or reference strains. Since the crystal structure of the C-terminal β-propeller domain of *P. aeruginosa* PilY1 (21) contains a bound Ca^2+^ ion, all the models with PilC/PilY1 included a Ca^2+^ ion. All the figures representing structural renderings were generated using ChimeraX (47). The protein-protein and protein-Ca^2+^ interfaces in the complexes have been analysed using ChimeraX (47) and/or PISA (48). The β-sheet motifs have been identified using PDBSum (49). Superposition of structures has been done using ChimeraX (47). FoldSeek (50) was used to search for structural homologs in the PDB (51) database. Signature protein domains have been identified by scanning the InterPro database (52). Protein domain distribution has been assessed by scanning UniProtKB (46). Prediction of signal peptides has been done using SignalP 6.0 (53) or manually for PilH proteins. Multiple sequence alignments have been performed using Clustal Ω (54). DNA Strider was used for routine analysis of protein sequences and computing their molecular weights (55).

### Bacterial strains and growth conditions

All the bacterial strains used in this study are listed in Table S3. *E. coli* DH5α strain was used for cloning and as a host for plasmid preparation. *E. coli* SHuffle T7 Express strain (New England Biolabs) was used for protein overexpression and purification. Bacteria were grown on solid or liquid lysogeny broth (LB) medium (BD Difco), or in a chemically defined medium (CDM) containing (50 mM NaPO_4_ pH 7.4, 0.05% NaCl). The CDM was supplemented with 0.4% glucose as a carbon source, 1 mM MgSO_4_, 100 µM CaCl_2_, 100 µM MnCl_2_, 50 µM ZnSO_4_, and 50 µM FeCl_3_. Depending on the isotopic labelling requirements of the experiment, the nitrogen source consisted of 1 g/l of either unlabelled NH_4_Cl or labelled ^15^NH_4_Cl. When required, LB or CDM liquid cultures were supplemented with 50 µg/mL kanamycin (Km), 150 µg/ml erythromycin (Ery), or 100 µg/ml ticarcillin.

The *N. meningitidis* strains used in this study are derivatives of a piliated and highly adhesive antigenic variant – sometimes called 2C43 – of the serogroup C clinical isolate 8013 (56). This variant has been previously sequenced, used to construct a complete collection of defined mutants (25, 35), and study pilus biogenesis in depth (19, 35). Meningococci were grown at 37°C in an atmosphere containing 5% CO_2_ on Gonococcal Base (GCB) agar (BD Difco) plates, supplemented with 12 μM FeSO_4_ and Kellogg’s supplements (57). For liquid cultures, we used GC-liquid medium [1.5% proteose peptone (BD Difco), 0.4% K_2_HPO_4_, 0.1% NaCl] when required with the addition of 12 μM FeSO_4_ and Kellogg’s supplements. Selection of *N. meningitidis* transformants was performed using 100 µg/ml Km, 2 µg/ml Ery, or 1 µg/ml nalidixic acid (Nal). Counter-selection was done using 20 mM *p*-chloro-phenylalanine (Cl-Phe) and 1% 2-deoxy-D-galactose (2-DOG).

Molecular biology experiments were performed according to standard procedures and/or the suppliers recommendations. QIAprep Spin Miniprep Kit (Qiagen) and Wizard Genomic DNA Purification Kit (Promega) were used for *E. coli* plasmid preparations and *N. meningitidis* genomic DNA extractions, respectively. Routine PCR was performed with GoTaq DNA polymerase (Promega). When high-fidelity PCR was required, we used Q5 High-Fidelity DNA Polymerase (New England Biolabs) or Pfu DNA Polymerase (Agilent Technologies). All the primers used in this study are listed in Table S4.

The pET-28b vector (Novagen) was used for protein overexpression in *E. coli*. All plasmids used in this study are listed in Table S5. To construct the plasmid for expressing 6His-PilK, we used as a template a plasmid carrying a synthetic *pilK* gene codon-optimised for expression in *E. coli* (synthesized by GeneArt). We amplified the portion of the gene encoding the soluble globular head of PilK using primers with *Nco*I and *Bam*HI overhangs, and cloned the PCR product into pET-28b cut by these restriction enzymes. This fused a non-cleavable N-terminal 6His tag to the N-terminus of truncated PilK. The same strategy was used to construct the plasmid for expressing 6His-PilK-PLUG, which has a C-terminally fused in-frame PLUG extension. This construction only necessitated a different reverse primer with a PLUG-encoding sequence. All the plasmids were verified by Sanger sequencing.

*N. meningitidis* mutants were constructed as follows. To construct markerless *ΔpilC1*, *ΔpilC2*, *ΔpilC1ΔpilC2* deletion mutants, we used a recently described dual counterselection strategy relying on the use of the *aphA3*-*pheS*-*galK* (APG) counterselection cassette (29). In the first step, we used overlap-extension PCR to generate PCR products in which upstream and downstream regions flanking *pilC1* or *pilC2* were spliced upstream and downstream the APG cassette. These PCR products were directly transformed in *N. meningitidis* allowing selection of *ΔpilC1*∷APG or *ΔpilC2*∷APG intermediates on Km plates. Meningococci were transformed as previously described (35). Bacteria were grown overnight (O/N) on GCB agar. The next day, bacterial cells were harvested and resuspended to an OD_600_ of 1 in GC-liquid medium supplemented with 12 μM FeSO_4_, Kellogg’s supplements, 2.5 mM MgCl_2_, and 2.5 mM MgSO_4_. Aliquots (300 µl) of the bacterial suspensions were mixed with 5 µl of PCR product and incubated for 30 min at 37°C, in 5% CO2 atmosphere, with shaking. Subsequently, 1 ml of supplemented GC-liquid medium was added, and the cultures were incubated during 3 h at 37°C. Transformants were recovered by plating the cultures onto GCB agar supplemented with Km. All the mutants were confirmed by PCR. In the second step, these intermediates were transformed directly, as above, with overlap-extension PCR products fusing the upstream and downstream regions flanking *pilC1* or *pilC2*. The markerless *ΔpilC1* or *ΔpilC2* mutants were obtained by applying Cl-Phe and 2-DOG counterselection against the APG cassette. The same two-step strategy was then applied to construct *ΔpilC1ΔpilC2* by deleting *pilC1* in the *ΔpilC2* mutant. However, since the *ΔpilC1*∷APG*ΔpilC2* intermediate is non-transformable, we needed to restore competence in the second step by adding 10 µg/ml of synthetic PLUG peptide. To construct triple mutants in the *ΔpilC1ΔpilC2* background, we used *pilK*, *pilQ*, *pilT* transposon (Tn) insertion mutants previously generated by *in vitro* transposition (25). All selected mutations were amplified by high-fidelity PCR using primers encompassing the Tn insertions and re-transformed in *ΔpilC1ΔpilC2* using 150 ng of PCR products and 10 µg/ml of synthetic PLUG peptide.

For genetic complementation of *ΔpilC* meningococcal mutants, we first cloned purified *pilC1* PCR products in the pMiniT vector (New England Biolabs) as described (58). To avoid phase variation, the poly(G) sequence present in the *pilC1* signal peptide was swapped by PCR mutagenesis by a sequence not prone to phase variation but without altering the protein sequence. The resulting plasmid was used as template to generate all the pNM99Ptet *pilC1* derivatives used for complementing *N. meningitidis* mutants (29). The plasmid carries within the *iga-trpB* intergenic locus from our parental strain, a P*tet* promoter that can be used to drive the expression genes of interest in the presence of 20 ng/ml aTC. Because this locus contains a DNA uptake sequence, the plasmid is efficiently taken up by natural transformation and integrates the P*tet*-driven gene by homologous recombination at the *iga-trpB* locus. To construct pNM99P*tet*-*pilC1*, the plasmid was linearised by PCR and *pilC1* was amplified using primers that introduce ends homologous to the termini of the linearised pNM99P*tet*. The two PCR products were fused by Gibson assembly. We generated, pNM99P*tet*-*pilK* and pNM99P*tet*-*pilK-PLUG* in a similar fashion. The pNM99P*tet*-*pilC1_Δplug_* plasmid was obtained by inverse PCR mutagenesis of pNM99P*tet*-*pilC1*, using a reverse primer designed to delete the sequence encoding the plug motif. This was followed by the recircularization of the resulting linear product to generate pNM99P*tet*-*pilC1_Δplug_*. Transformations were done with 150 ng of plasmid and transformants were selected on GCB agar supplemented with Ery and verified by PCR. When complementing non-competent *ΔpilC1ΔpilC2* and *ΔpilC1ΔpilC2pilK* mutants, the PLUG peptide was supplemented at 10 µg/ml throughout all the procedure.

### Quantifying meningococcal competence

To determine transformation frequencies, bacterial colonies were resuspended in GC-liquid medium supplemented with 5 mM MgSO_4_ and adjusted to an OD_600_ of 0.1. One µg of genomic DNA carrying a cassette promoting resistance to Nal was added to 200 µl of the bacterial suspension in the presence or absence of 10 µg/ml synthetic peptides, and incubated for 20 min at 37°C in 5% CO_2_ atmosphere with shaking. Then, 800 µl of transformation liquid medium (containing 12 μM FeSO_4_ and Kellogg’s supplements, except L-Glutamine) was added as described (59), and bacteria were incubated without agitation for 2 h at 37°C. Bacterial serial dilutions were then plated on agar plates with and without Nal, incubated O/N at 37°C, and CFU counts were performed the next day. The transformation frequencies were calculated by dividing the number of CFU obtained on Nal-containing agar plates by the number of CFU obtained on antibiotic-free agar plates. If no colonies were obtained at the lowest dilution, a CFU of 0.9 was arbitrarily assigned and the point was marked as below the detection limit.

### Quantifying meningococcal piliation

This was done by performing immunofluorescence assays on bacteria from the above transformation, after the 2 h incubation step at 37°C without agitation. In brief, 2 x 10⁷ bacteria were seeded onto coverslips coated with 1 mg/ml poly-D-lysine and placed in the wells of a 24-well plate. Following centrifugation at 2,500 *g* for 5 min to promote bacterial adhesion onto the coverslips, the plate was incubated at 37°C for 40 min. The supernatant was then carefully aspirated, and the wells were rinsed once with PBS. Bacteria were fixed by incubation with 500 µl of 4% paraformaldehyde (PFA) in PBS for 20 min. After fixation, the wells were washed with PBS, and the coverslips were processed for immunostaining. The coverslips were washed with 50 mM NH_4_Cl (in PBS) for 5 min, then blocked with 0.1% BSA (in PBS) for 30 min. Pili were stained with the 20D9 mouse monoclonal antibody that specifically recognises the filaments of our parental strain (60), used at 1/1,000 dilution. A goat anti-mouse antibody conjugated with Alexa Fluor 546 (Thermo Fisher Scientific) was used as the secondary antibody at 1/400 dilution. Bacteria were detected by DNA staining using 100 ng/ml 4’,6-Diamidino-2-phenylindole dihydrochloride (DAPI). Coverslips were mounted in Mowiol before images were captured using a spinning disk confocal CrestOptics X-Light V3 microscope. The images were processed using the VisiView software (Visitron Systems). Piliation indexes were calculated by image analysis as the ratio of the areas occupied by the fluorescent signals for pili and DNA, respectively. Because the DNA signal is proportional to the number of bacteria, the above ratio provides a relative measurement of piliation normalised to cell number in arbitrary units. All image analyses were performed using the FIJI software (61). Statistical analyses were performed with Prism (GraphPad Software). Comparisons were done by one-way ANOVA, followed by Dunnett’s multiple comparison tests. An adjusted *P* < 0.05 was considered significant (\**P* < 0.05, \*\**P* < 0.01, \*\*\**P* < 0.001, \*\*\*\**P* < 0.0001).

### 3FLAG-PLUG immunofluorescence staining

To detect the 3FLAG-PLUG by immunostaining, bacteria were resuspended in 5 ml DMEM (Gibco) supplemented with 10% foetal bovine serum (Gibco) to an OD_600_ of 0.1 and incubated in the presence of 40 µg/ml synthetic peptide for 2 h at 37°C, in 5% CO_2_ atmosphere, with shaking. Then, 2 x 10⁷ bacteria were seeded onto coverslips, fixed and blocked as above. Prior to blocking, cells were permeabilized for 10 min with 0.1% Triton X-100 and 0.2 mg/ml lysozyme. The 3FLAG-PLUG peptide was stained with a monoclonal mouse anti-FLAG-tag antibody (Sigma), diluted 1/1,000 in PBS containing 0.1% BSA. This was followed by incubation with an Alexa Fluor 546-conjugated goat anti-mouse secondary antibody (Thermo Fisher Scientific), used at 1/400 dilution. Bacterial DNA was counterstained with DAPI (100 ng/ml). After washing and mounting, immunofluorescence images were acquired on an SP8 laser scanning confocal microscope (Leica Microsystems) and deconvolved using Huygens software (Scientific Volume Imaging).

### Protein purification

To purify ^15^N-labelled 6His-PilK or 6His-PilK-PLUG proteins for NMR analysis, the corresponding pET-28b derivatives were transformed into *E. coli* SHuffle T7 Express and grown on LB plates O/N. A single colony was used to inoculate a starter culture in LB supplemented with Km for approximately 8 h, then diluted 1/50 into CDM containing Km and ^15^NH_4_Cl (1 g/l), and grown O/N at 37°C. The following morning, the culture was back diluted to OD_600_ 0.05 in 1 l of the same medium. Once the OD_600_ reached 0.2, the cultures were cooled down to 16°C, and protein expression was induced O/N with 1 mM isopropyl 1-thiogalactopyranoside (IPTG) (Merck Chemicals). The next day, the cells were harvested by centrifugation at 8,000 *g* for 15 min and the pellets were frozen at -80°C. The cell pellet was resuspended in 15 ml of binding buffer (50 mM NaPO_4_ pH 8.0, 300 mM NaCl, 20 mM imidazole) supplemented with complete EDTA-free Protease Inhibitor Cocktail (Sigma), 20 ng/ml DNase I, 1 mM MgCl_2_ and 1 mM phenylmethylsulphonyl fluoride, and lysed using a French press. The lysate was then clarified by centrifugation at 11,000 *g* for 30 min.

6His-PilK and 6His-PilK-PLUG proteins were purified by immobilised metal affinity chromatography on an ÄKTA go using 1 ml HisTrap HP columns (Cytiva) according to the manufacturer’s instructions. After washing with binding buffer (50 mM NaPO_4_ pH 8.0, 300 mM NaCl, 20 mM imidazole), proteins were eluted by applying an imidazole gradient up to 300 mM. Sample purity was assessed by SDS-PAGE and Coomassie staining. Fractions containing the protein of interest were pooled and further purified by size-exclusion chromatography (SEC) on a Superdex 75 Increase 10/300 GL column (Cytiva) equilibrated with 50 mM NaPO_4_ pH 6.5, 100 mM NaCl and 3% glycerol. Protein concentrations were determined spectrophotometrically using a NanoDrop Lite (Thermo Fisher Scientific).

To purify unlabelled 6His-PilK and 6His-PilK-PLUG proteins for crystallography, BLI and Thermofluor assays, the protein expression and purification were carried out as described above, with CDM medium supplemented with unlabelled NH_4_Cl. The proteins were affinity-purified using Gravity Flow Chromatography columns (Bio-Rad) loaded with 1.5 ml Ni-NTA agarose resin (Qiagen) preequilibrated with binding buffer (50 mM NaPO_4_ pH 8.0, 300 mM NaCl, 20 mM imidazole). The resin was mixed with the clarified lysates and the columns were allowed to drain. The columns were then washed with binding buffer several times, before the proteins were eluted with elution buffer (50 mM NaPO_4_ pH 8.0, 300 mM NaCl, 300 mM imidazole). Both proteins were further purified by SEC on an ÄKTA go using a Superdex 75 16/600 GL column (GE Healthcare), and simultaneously buffer-exchanged into crystallization buffer (10 mM HEPES pH 7.5, 150 mM NaCl) or BLI/Thermofluor buffer (10 mM HEPES pH 7.5, 100 mM NaCl).

### Quantifying protein-protein interactions by bio-layer interferometry

Quantification of the interaction between 6His-PilK and various synthetic peptides was performed by BLI using the Octet R8e instrument (Sartorius) as described (62). Streptavidin (SA) biosensors (Sartorius) were hydrated for 15 min in kinetic buffer [PBS, 0.1% (w/v) BSA, 0.02% (v/v) Tween 20]. For the binding of 6His-PilK to the biosensors, the purified protein was biotinylated as follows. It was first buffer-exchanged in 1X PBS using a PD-10 desalting column (Cytiva) and biotinylated using NHS-PEG4-Biotin Ester (Thermo Scientific) as directed by the manufacturer. Specifically, 6His-PilK at 1 mg/ml was incubated with 1 mM NHS-PEG4-Biotin Ester for 30 min at room temperature, at a 1:3 molar ratio protein to biotin. Following the reaction, the biotin in excess was removed using a 7 kDa Zeba spin desalting column (Fisher Scientific).

In parallel, synthetic peptides used as analytes were resuspended in 1X PBS at 6.5, 50, 200 µM (PLUG peptide) or only at 50 µM (GLUP and PLUG+/-peptides), and placed into black-bottom 384-well microplates (Greiner Bio-One). After 6His-PilK loading at a concentration of 10 µg/ml, a baseline step was conducted to remove any unbound protein before the association step, during which the biosensors were immersed in the wells containing the peptides. Finally, a dissociation step was performed by transferring the biosensors back into the kinetic buffer. For each interaction assay, a reference biosensor without immobilised 6His-PilK was included in parallel to account for nonspecific binding of the peptides, and the corresponding signal was subtracted. All the measurements were taken at 25°C with shaking at 1,000 rpm. The data were analysed using Sartorius Octet Research Suite Software Version 14.

### Determining protein structure by X-ray crystallography

Crystallization trials with 6His-PilK-PLUG at 22 mg/ml in (10 mM HEPES pH 7.5, 150 mM NaCl) were performed by the sitting-drop vapour-diffusion method in SwissCi 96-well crystallization plates (STPLabTech), using a Mosquito crystallization robot (STPLabTech) and a series of commercial screens. The first crystal hits emerged from a condition of the Nextal PEGs Suite (Calibre Scientific) containing 25% (v/v) PEG550 mono-ethyl ether and 0.1 M Na-acetate buffer at pH 4.5. After several optimisation rounds, well-diffracting crystals were obtained by the sitting-drop vapour-diffusion method by mixing 200 nl protein solution with 200 nl reservoir solution composed of 16.4% PEG550 mono-ethyl ether and 0.1 M Na-acetate buffer at pH 5.72. Crystals were cryo-protected with reservoir solution supplemented with 25% (v/v) glycerol prior flash-cooling in liquid nitrogen. X-ray diffraction data were acquired at beam line ID30B at the European Synchrotron Radiation Facility in Grenoble. Diffraction data were reduced with the xia2 pipeline (63) and scalded and merged using the CCP4 (64) suite of programs POINTLESS (65), AIMLESS (66) and TRUNCATE (67).

The structure of 6His-PilK-PLUG was solved by molecular replacement with the program *Phaser* (68) using an AlphaFold 3 prediction as the search model. The crystals belong to space group *P*2_1_, with six molecules per asymmetric unit (Fig. S3). The protein chains were corrected automatically with the program Buccaneer (69). Refinement and model adjustment were carried out with the programs REFMAC5 (70) and Coot (71), respectively. A random set of 4.8% of reflections was set aside for cross-validation purposes. For maximum-likelihood refinement with REFMAC5 TLS refinement and local NCS restraints were implemented. Model quality was assessed with internal modules of Coot (71) and using the MolProbity server (72). Atomic coordinates and structure factors have been deposited within the PDB (51) with accession number 32YF. Data collection and refinement statistic are provided in Table S6.

### Testing protein stability by performing Thermofluor assays

Thermal stability assays of 6His-PilK or 6His-PilK-PLUG have been done as described (34), with the following modifications. Purified proteins at 10 µM were incubated in the presence of 20X SYPRO Orange (Sigma-Aldrich) in a total volume of 20 µl. Samples were then heated using a Bio-Rad CFX96 Touch Real-Time PCR instrument from 25 to 85°C at a rate of 0.5°C per 30 sec. Protein unfolding was detected by monitoring changes in SYPRO Orange fluorescence. The Tm were determined using the first derivative values of raw fluorescence data using Bio-Rad CFX96 Manager version 3.1 software.

### Testing protein stability by NMR spectroscopy

Purified protein samples were concentrated to 250 µM in NMR buffer (50 mM NaPO_4_ pH 6.5, 100 mM NaCl, 5% DMSO, 10% D_2_O) then loaded into 5 mm NMR tubes. ^1^H-^15^N HSQC spectra were collected, at the specified temperatures and timeframes, on a Bruker Avance NEO 600 MHz spectrometer equipped with a triple-resonance cryogenic probe. Data were processed using TopSpin 4 (Bruker) and visualised in CcpNmr version 3 (73).

## ACKNOWLEDGEMENTS

This work was supported by the Agence Nationale de la Recherche: ANR-24-CE11-3366 (T4Ptip) and ANR-10-INBS-0005 (FRISBI). We acknowledge the European Synchrotron Radiation Facility (ESRF) for provision of synchrotron radiation on beamline ID30B under proposal number MX-2799 (74). We are grateful to Emilia Mauriello (NanoBacVir), Xavier Nassif (INEM) and Mike Koomey (University of Oslo) for critical reading of this manuscript. We thank Christophe Piesse from the peptide biosynthesis platform (Institut de Biologie Paris-Seine) for the synthesis of peptides, Olivier Bornet from the NMR platform (Institut de Microbiologie de la Méditerranée) for help with the spectrometer, Yann Denis from the Transcriptome platform (Institut de Microbiologie de la Méditerranée) for help with the Real-Time PCR instrument, Alain Roussel (LISM) for help with the BLI experiments, and the ESRF staff for assistance during X-ray diffraction experiments.

## DATA AVAILABILITY

The crystal structure of the chimeric PilK-PLUG protein from N. meningitidis has been deposited at the PDB with accession code <u>32YF</u>. All the data generated during this study are included in this paper and/or its Supplementary information file. Source data are provided with this paper.

## AUTHORS CONTRIBUTION

VP was responsible for conception of the work. NB, LE, GS, MC and VP contributed to the design and supervision of the experiments. MD, MW, VRZ and OV contributed to the acquisition and analysis of data. VP drafted the article. MD, MW, NB, LE, GS and MC revised the article critically. All authors approved the final version.

